# Secretory Leukocyte Protease Inhibitor Protects Against Severe Urinary Tract Infection in Mice

**DOI:** 10.1101/2023.10.10.561753

**Authors:** Anne L. Rosen, Michael A. Lint, Dayne H. Voelker, Nicole M. Gilbert, Christopher P. Tomera, Jesús Santiago-Borges, Meghan A. Wallace, Thomas J. Hannan, Carey-Ann D. Burnham, Scott J. Hultgren, Andrew L. Kau

**Author notes:** Dayne H. Voelker; Mayo Clinic, 200 First Street SW, Rochester, MN Meghan A. Wallace; Pattern Bioscience, 9600 Great Hills Trail, Ste 160E, Austin, TX.

## Abstract

Millions suffer from urinary tract infections (UTIs) worldwide every year with women accounting for the majority of cases. Uropathogenic *Escherichia coli* (UPEC) causes most of these primary infections and leads to 25% becoming recurrent or chronic. To repel invading pathogens, the urinary tract mounts a vigorous innate immune response that includes the secretion of antimicrobial peptides (AMPs), rapid recruitment of phagocytes and exfoliation of superficial umbrella cells. Here, we investigate secretory leukocyte protease inhibitor (SLPI), an AMP with antiprotease, antimicrobial and immunomodulatory functions, known to play protective roles at other mucosal sites, but not well characterized in UTIs. Using a mouse model of UPEC-caused UTI, we show that urine SLPI increases in infected mice and that SLPI is localized to bladder epithelial cells. UPEC infected SLPI-deficient (*Slpi^-/-^*) mice suffer from higher urine bacterial burdens, prolonged bladder inflammation, and elevated urine neutrophil elastase (NE) levels compared to wild-type (*Slpi^+/+^*) controls. Combined with bulk bladder RNA sequencing, our data indicate that *Slpi^-/-^* mice have a dysregulated immune and tissue repair response following UTI. We also measure SLPI in urine samples from a small group of female subjects 18-49 years old and find that SLPI tends to be higher in the presence of a uropathogen, except in patients with history of recent or recurrent UTI (rUTI), suggesting a dysregulation of SLPI expression in these women. Taken together, our findings show SLPI protects against acute UTI in mice and provides preliminary evidence that SLPI is likewise regulated in response to uropathogen exposure in women.

## Importance

Annually, millions of people suffer from urinary tract infections (UTIs) and more than $3 billion are spent on work absences and treatment of these patients. While the early response to UTI is known to be important in combating urinary pathogens, knowledge of host factors that help curb infection is still limited. Here, we use a mouse model of UTI to study secretory leukocyte protease inhibitor (SLPI), an antimicrobial protein, to determine how it protects the bladder against infection. We find that SLPI is increased during UTI, accelerates the clearance of bacteriuria, and upregulates genes and pathways needed to fight an infection while preventing prolonged bladder inflammation. In a small clinical study, we show SLPI is readily detectable in human urine and is associated with the presence of a uropathogen in patients without a previous history of UTI, suggesting SLPI may play an important role in protecting from bacterial cystitis.

## Introduction

In the United States, urinary tract infections (UTIs) lead to over 10 million office visits and cost more than $3 billion annually in sick leave and treatment (1,2). Women are at higher risk of developing frequent UTIs than men (3–5) and the primary culprit of UTIs is uropathogenic *Escherichia coli* (UPEC), a gram-negative bacterium responsible for most community-acquired infections (6,7). UPEC strains are common gut commensals and do not cause disease in the gastrointestinal (GI) tract, but once shed, they can colonize the periurethral tissues (6,8). UPEC is known to utilize multiple virulence factors to aid its ascent, adherence, and invasion of the bladder epithelium (9). Among these virulence factors, type 1 pili, a hair-like extracellular structure, are essential for adhesion and invasion of UPEC to the luminal superficial facet epithelial cells of the bladder (7,10,11). Once inside the bladder epithelium, UPEC can escape the immune response by replicating within the facet cells to form intracellular bacterial communities (IBCs) that evade phagocytosis by neutrophils (12–14). In response to infection, superficial epithelial cells undergo a process similar to apoptosis and are exfoliated into the urine (10,15).

While the adaptive immune system responds to UPEC-caused UTI, the innate immune response is thought to play a key role in controlling infection (16). Toll-like receptors (TLR) on host cells recognize conserved molecules like LPS (17) and flagellin (18) expressed on the surface of gram-negative bacteria like UPEC. Once TLR4 recognizes LPS, the NF-κB pathway becomes activated resulting in the transcription of inflammatory cytokines including IL-1β, IL-6, and TNFα (19,20). Accumulation of these molecules leads to the recruitment of neutrophils which is crucial in controlling early infection with UPEC (20–22). However, recruitment of neutrophils is carefully regulated to maintain a balance necessary to clear infection while protecting host tissues. An inadequate number of neutrophils can result in failure to clear UPEC from the bladder, but an overabundance of neutrophils can lead to chronic cystitis and excessive tissue damage (23,24), emphasizing the delicate homeostasis that must be maintained to resolve UTI.

Increased levels of proinflammatory cytokines also upregulate expression of antimicrobial peptides (AMPs), which help prevent invasive bacterial infections (25,26). AMPs are small, positively charged proteins that are commonly produced by neutrophils, macrophages, and epithelial cells as well as other cell types. While they can be constitutively expressed, their production is also induced by microbial stimuli or tissue damage (27,28). While AMPs including LL-37 (29,30) and serpins (24), have been studied in the context of UTI, little is known about the roles and functions of the well characterized AMP secretory leukocyte protease inhibitor (SLPI) which is also found in the urogenital tract (31,32). SLPI has been extensively studied in the respiratory tract (33), but has also been shown to help combat urogenital infections from HIV, gonorrhea, and *Candida* yeast (34–36), though its function in bacterial UTIs remains unknown. SLPI is a small, cationic protein containing two domains, each with their own respective functions (37–40). The N-terminal domain is associated with antimicrobial activity and has been reported to demonstrate bactericidal activity against *E. coli* and *S. aureus in vitro* (38). The C-terminal domain broadly inhibits serine proteases and may be important in limiting inflammation caused by enzymes, including those released by activated neutrophils such as cathepsin G and neutrophil elastase (NE) (24,39,41). NE is a serine protease stored in preformed granules (42) where it aids the immune system by digesting phagocytosed proteins (43). NE released from activated neutrophils can also degrade extracellular matrix proteins like elastin (43,44) and fibronectin (45) resulting in host tissue damage (43,45–47). Prior reports show SLPI can directly inhibit NE protease function and reduce damage in response to inflammation (28,48). Additionally, SLPI has been shown to regulate the immune response through the direct blocking of the NF-κB promoter and by preventing the degradation of NF-κB inhibitors (49–51). Other serine protease inhibitors from the serpin family are also increased during UTI (24).

In this study, we tested the hypothesis that SLPI protects the bladder from UPEC-caused cystitis using a mouse model of UTI. We found that mice respond to the introduction of UPEC in the urinary tract by upregulating SLPI within hours of infection. We also found that *Slpi^-/-^* mice demonstrate evidence of a dysregulated immune response to UPEC infection, while also exhibiting an increased burden of UPEC in the urine and delayed resolution of inflammation at later time points. We complement these findings in mice with a small clinical study showing that women with acute bacteriuria tend to have elevated SLPI levels, providing preliminary evidence that SLPI plays a protective role human UTIs.

## Results

### SLPI is increased in the mouse urinary tract following UTI

We used an established mouse model of UTI to determine whether SLPI levels are increased in the urine following infection (52). Briefly, we grew a kanamycin-marked strain of UTI89, a prototypical UPEC clinical isolate (12), under static conditions to induce expression of type 1 pili (11,53) before transurethrally inoculating ∼1 x 10^8^ CFU or sterile PBS directly into the bladders of C57BL/6J female mice. We found the concentration of SLPI in urine was significantly increased in infected mice compared to mock-infected mice as early as 3 hours post infection (hpi) and that this concentration peaked at 7 hpi before slowly decreasing over a four-day period (Figure 1A). A time series ANOVA showed that infected mice had significantly higher levels of SLPI over the course of infection compared to mock-infected controls (Treatment, p=1.5×10^-6^). Additionally, transcription of *Slpi* was increased in bladder homogenates 7 hpi, suggesting that SLPI protein found in the urine is originating, at least in part, from the bladder and not solely as increased urinary secretion of SLPI from the kidneys (Figure 1B). Interestingly, bacteriuria (UPEC in the urine) peaked 1 day after infection (Figure 1C), long after urine SLPI levels reached their apex, suggesting that SLPI is involved in the early innate response to infection. To determine how well urine SLPI levels correlated to bacteriuria, we created a receiver operating characteristic (ROC) curve for each timepoint. These ROC curves show that the amount of SLPI is correlated to bacteriuria in mice at each time point we examined from 3 hpi to 4 days post infection (dpi) with an AUC value of 0.79 or greater (Figure 1D).

**Figure 1:**
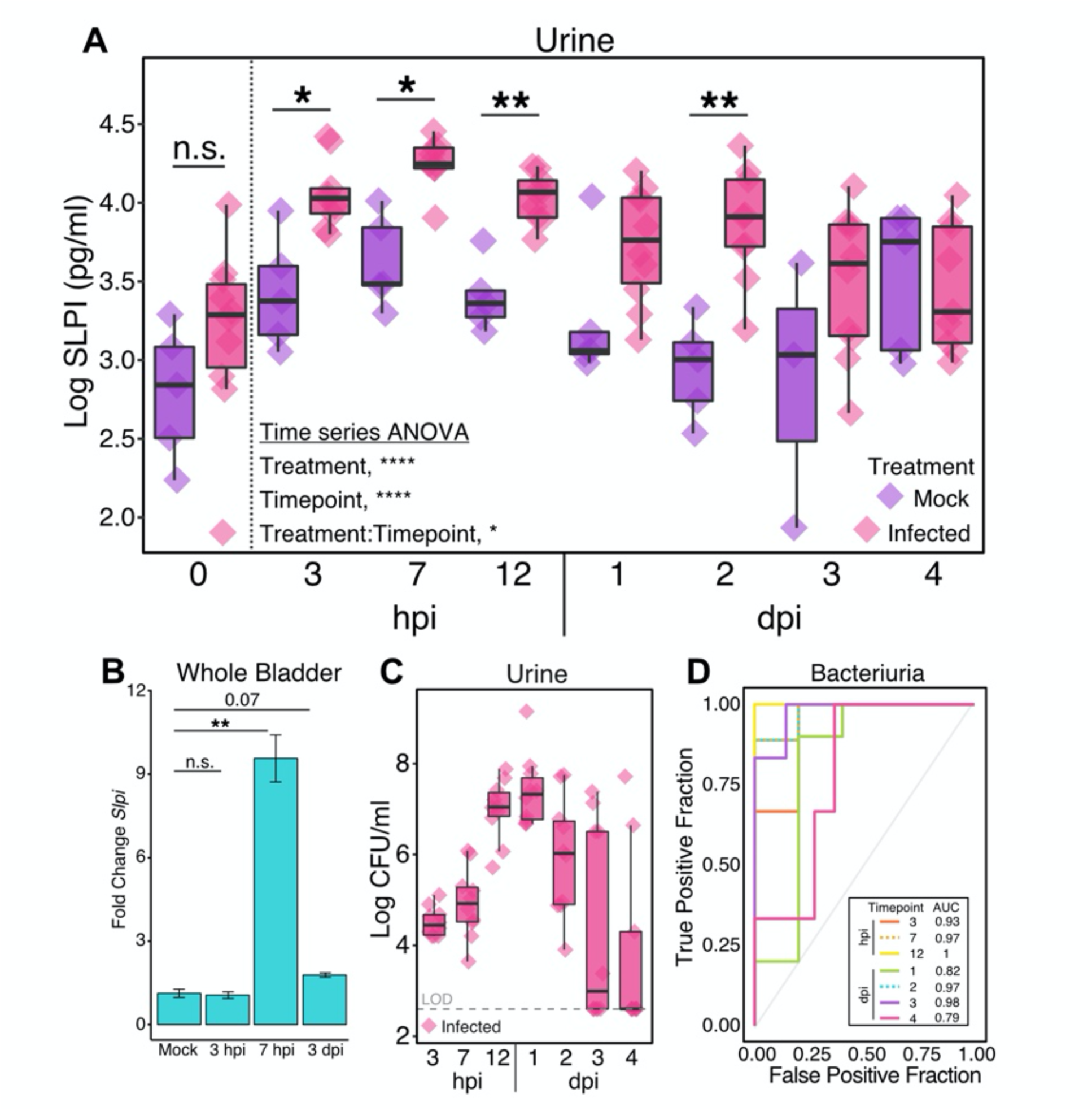
SLPI is increased in mouse urinary tract following UTI. **A)** Urine SLPI levels in C57BL/6J mock-infected (n=3-5) and UTI89-infected (n=9-10) mice. A time series ANOVA was performed on all timepoints after 0 including imputed values for missing samples (2 mock and 5 infected). A post-hoc Student’s t-test was performed on imputed data with FDR correction. **B)** qRT-PCR of *Slpi* in whole bladder homogenates at 3 hpi (n=12), 7 hpi (n=8), and 3 dpi (n=3) represented as fold change to mock-infected mice (n=3-4) (Student’s t-test). **C)** Log CFU/ml of UTI89 in urine of infected mice over time. Dashed line represents limit of detection (LOD). **D)** ROC curve analysis using SLPI levels in (A) to classify bacteriuria shown in (**C**) with area under the curve (AUC) values listed for each timepoint.

### SLPI is expressed in the bladder epithelium

To better understand localization of SLPI expression in the urinary tract, we collected bladders from *Slpi^+/+^* and *Slpi^-/-^*mice 7 hours after infection and performed immunofluorescent staining against SLPI protein (Figure 2) along with isotype controls (Figure S1). Staining with a SLPI-specific antibody revealed a strong signal on the luminal surface of the bladder, consistent with staining of the uroepithelium in both the mock-infected and UTI89-infected mice (Figure 2A and 2B). Control staining with an isotype antibody (Figure S1A-C) or against *Slpi^-/-^* mouse bladders (Figure S1D) failed to produce noticeable signal, indicating that the observed staining was specific to SLPI. While we did observe some staining of SLPI in the tissues underlying the epithelium, it was less prominent than the signal observed in the uroepithelium, implying that SLPI is primarily expressed by the epithelium in the bladder. Next, we quantified the difference in SLPI fluorescence of the uroepithelium between mock and infected mice. While there was no significant difference in DNA fluorescence (Figure 2C), we did see a decreased SLPI signal in infected mice both before (Figure 2D) and after normalizing to DNA fluorescence (Figure 2E). This finding implies that SLPI protein is largely depleted from the bladder epithelium 7 hpi either through exfoliation of bladder epithelial cells or secretion, while transcription of SLPI is upregulated (Figure 1B). To better understand the dynamics between SLPI transcription and secretion, we examined the human bladder epithelial cell line, 5637, to determine the epithelial response of SLPI to UPEC exposure. We found that infection with UTI89 resulted in significantly increased secretion of SLPI protein into the culture supernatant by two hours of infection compared to cells exposed to PBS alone (Figure 2F). However, the increases in SLPI protein apparent in the culture supernatant were not accompanied by increases in cellular transcripts of SLPI at that timepoint (Figure 2G), suggesting that increases in SLPI quantity in the urinary tract may not be transcriptionally mediated in the period immediately after UTI89 exposure.

**Figure 2:**
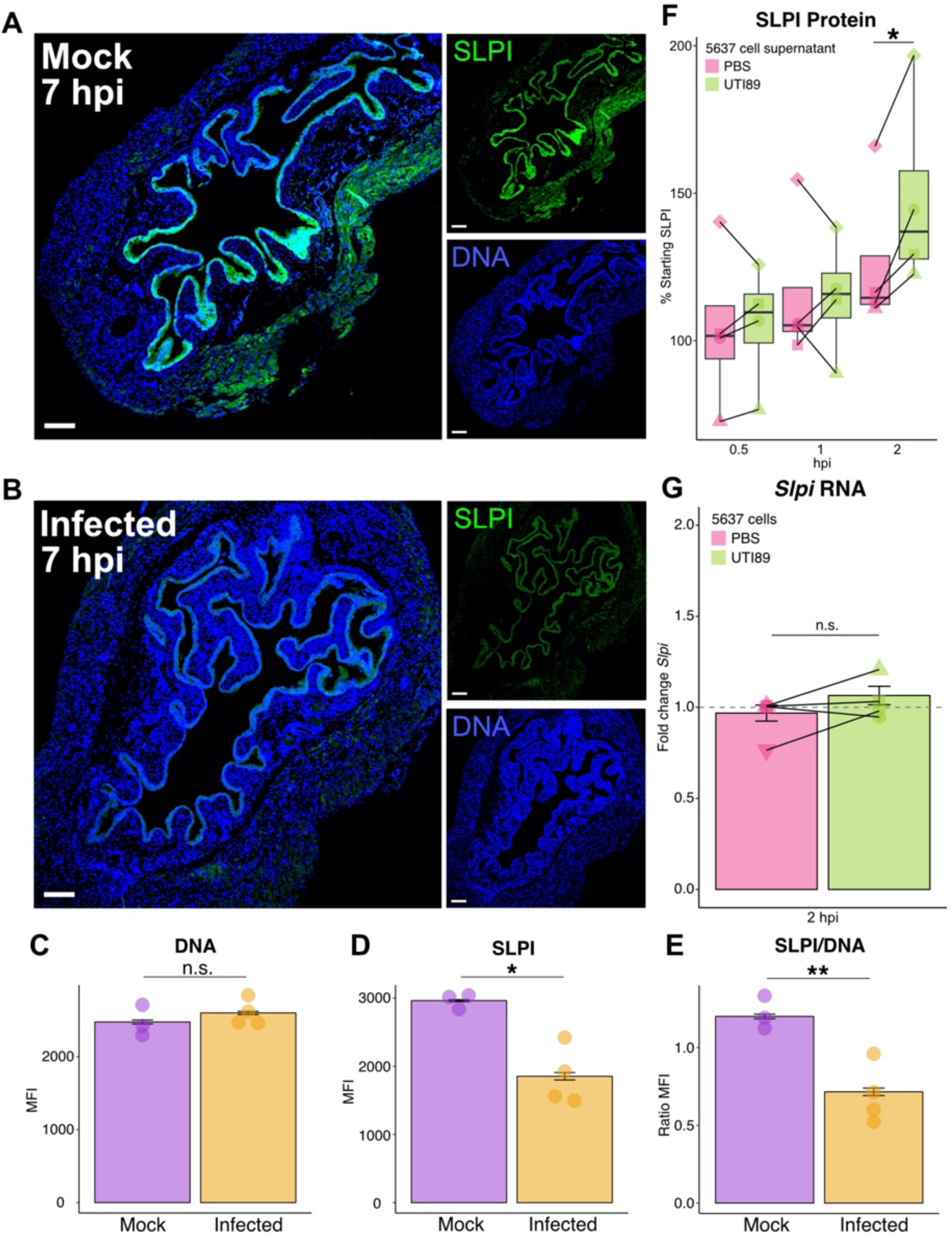
SLPI is expressed in the bladder epithelium. **A-B)** Immunofluorescence staining of SLPI protein expression (green) and DNA using Hoechst stain (blue) in mock-infected (**A**) and infected (**B**) bladders from *Slpi^+/+^* mice. Left panels (5X magnification, 200 μm scalebars) are merged images of SLPI and DNA shown as smaller panels on the right. **C-E)** Mean fluorescence intensity (MFI) of mock and infected bladder epithelium for DNA (C) and SLPI (D) with MFI ratio of SLPI/DNA (E) (Student’s t-test). **F)** Comparison of supernatant SLPI levels in control (PBS) or infected (UTI89) 5637 bladder epithelial cells represented as percent of starting SLPI in each well. **G)** Fold-change in *Slpi* transcription as measured by qRT-PCR in 5637 cells at 2 hpi. For (F) and (G), shapes denote separate biological replicate experiments, each representing the average of 4 or more wells (Paired Student’s t-test).

### *Slpi^-/-^* mice have prolonged bacteriuria compared to *Slpi^+/+^* mice

To investigate whether SLPI is involved in susceptibility to UPEC-caused UTI, we utilized 129;BL/6 SLPI^tm1Smw^/J mice (28), which are engineered to lack SLPI, and back-crossed them onto a C57BL/6J background for at least 10 generations (*Slpi^-/-^* mice). After infecting mice with UTI89, we found that compared to wild-type (*Slpi^+/+^*) mice, *Slpi^-/-^* mice had significantly more bacteria in their urine at 2 and 3 dpi (Figure 3A). We also measured the abundance of neutrophils in urine sediments (pyuria) following infection with UTI89 and found that they were significantly increased in *Slpi^-/-^* mice compared to *Slpi^+/+^* mice 7 hours after infection but returned to *Slpi^+/+^* levels by 24 hpi (Figure 3B). The increased urine neutrophils did not correspond to differences in urine titers at this timepoint (Figure 3A) suggesting that a lack of SLPI may lead to more pronounced early neutrophil recruitment that does not impact bacterial titers. We also did not see any significant differences in bladder or kidney bacterial burdens over the 7 day UTI model (Figure 3C and 3D; see also Figure S2A-C). We next used an *ex vivo* gentamicin protection assay (54) to test whether UTI89 invasion into bladder cells was altered in *Slpi^-/-^* mice but found no significant differences from *Slpi^+/+^* (Figure 3E). Taken together, we observed that the timepoint where urine SLPI is first increased does not coincide with reduced UTI89 urine titers, suggesting that SLPI is not directly inhibiting UTI89 growth in our model.

**Figure 3:**
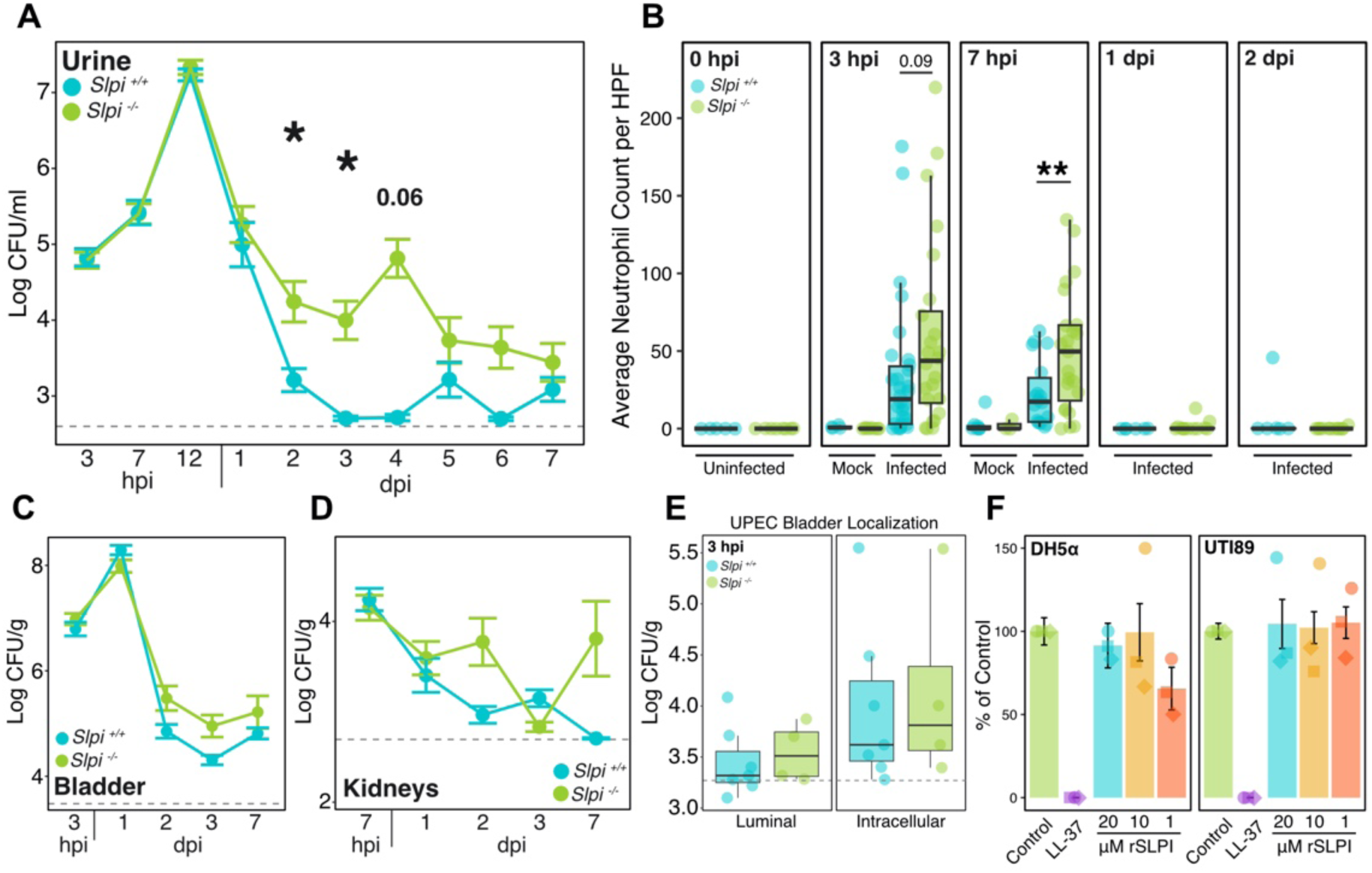
*Slpi^-/-^* mice have prolonged bacteriuria compared to *Slpi^+/+^* mice. **A)** Urine bacterial titers from of *Slpi^+/+^* (blue) and *Slpi^-/-^* mice (green) during one week of infection with UTI89. Data are log base 10 transformed CFU/ml and are combined from 7 experiments with 5-37 mice per genotype per timepoint (Student’s t-test). **B)** Averaged cytospin cell counts of neutrophils in urine from uninfected (n=5-8), mock-infected (n=3-8) and UTI89-infected (n=6-26) *Slpi^+/+^* (blue) and *Slpi^-/-^*mice (green) from 0 hpi to 2 dpi. Data is combined from 4 separate experiments (Student’s t-test). **C)** Bladder bacterial titers in log CFU/g. Data are combined from 4 experiments with 5-14 mice per genotype per timepoint. **D)** Kidney bacterial titers in log CFU/g. Data are combined from 7 experiments with 5-19 mice per genotype per timepoint. **E)** *Ex vivo* gentamicin protection assay on infected bladders from *Slpi^+/+^* (blue) and *Slpi^-/-^* mice (green) showing log CFU/g of tissue for recovered UTI89 from luminal or intracellular samples. For (A) – (E), gray dashed lines on each plot represent average limit of detection for that sample type. **F)** *In vitro* recombinant SLPI (rSLPI) antimicrobial activity against UTI89 and DH5α in PBS + 1% LB. All samples were normalized to PBS control group (green) and 1 uM LL-37 antimicrobial protein was used as a positive control (purple). The final concentrations of rSLPI in each culture were: 20 uM (blue), 10 uM (yellow), and 1 uM (orange).

Next, we wanted to test if SLPI could inhibit the growth of UTI89 *in vitro*. Using an approach based on previously described protocols (38,55), we tested if exposure to a commercially available recombinant SLPI (rSLPI) protein could reduce the viability of UTI89. We found that even after supraphysiological exposures to rSLPI, the presence of rSLPI did not significantly reduce the viability of UTI89 or a lab strain of *E. coli* (DH5a) (Figure 3F). In contrast, exposure to LL-37, which is known to kill *E. coli* (56), readily reduced the viability of both UTI89 and DH5a in the same assay. These results show that SLPI does not exert strong antimicrobial activity against *E. coli* and supports the idea that other functions of SLPI likely mediate its effect on UTI in our mouse model.

### *Slpi^-/-^* mice demonstrate a dysregulated immune response in the bladder at 1 dpi

To identify host responses that account for the differences in urine UPEC titers between *Slpi^+/+^* and *Slpi^-/-^* mice, we performed bulk RNA sequencing on whole bladders. We included four groups of mice: (1) *Slpi^+/+^* mice undergoing mock-infection (M-*wt*); (2) *Slpi^+/+^* mice infected with UTI89 (I-*wt*); (3) *Slpi^-/-^*mice undergoing mock-infection (M-*ko*); and (4) *Slpi^-/-^* mice infected with UTI89 (I-*ko*). Mice were sacrificed and bladder tissues harvested for RNAseq 1 day after inoculation (Figure 4A).

**Figure 4:**
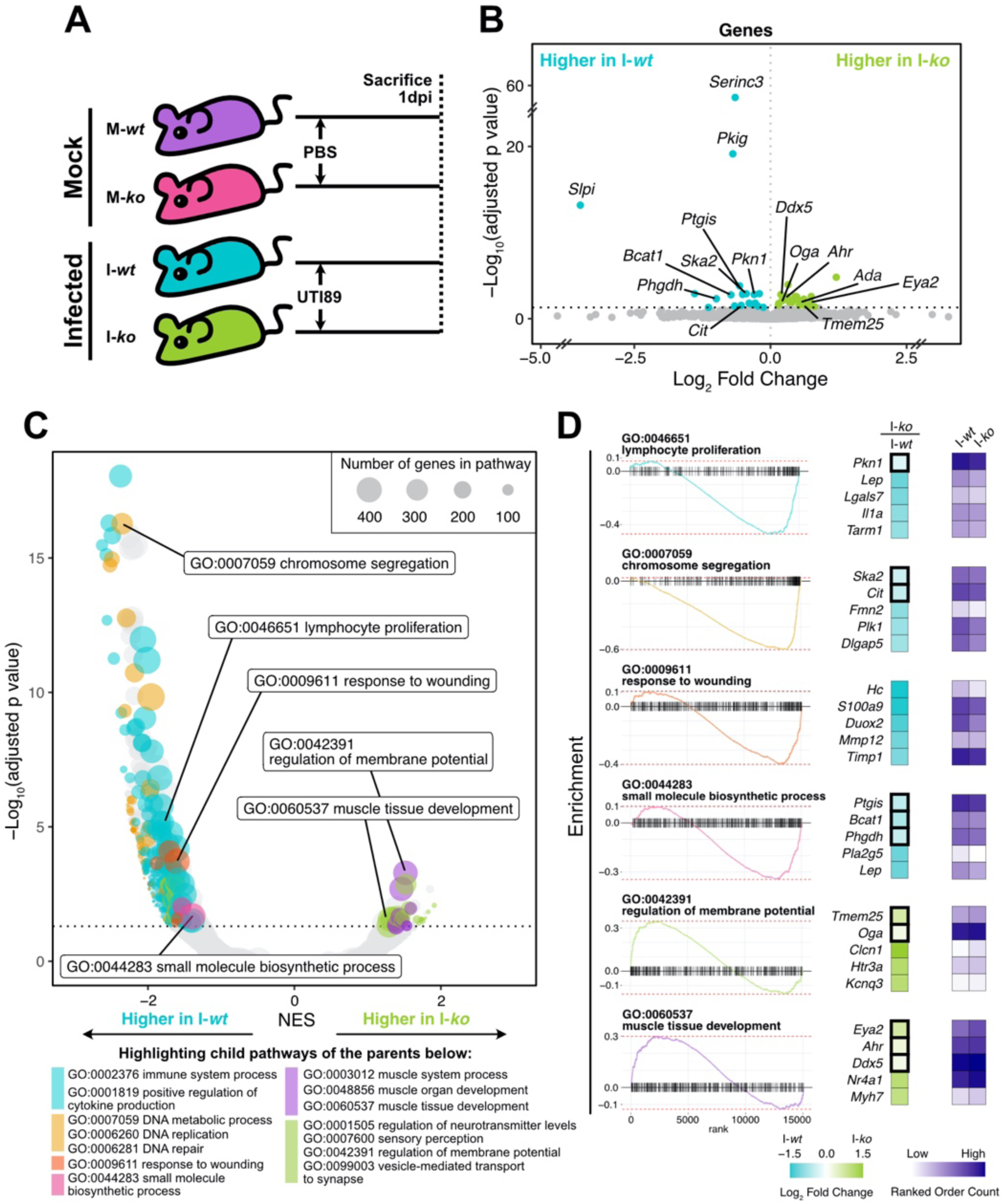
*Slpi^-/-^* mice demonstrate a dysregulated immune response in the bladder at 1 dpi. **A)** Schematic of experimental approach to assess differences in whole bladder transcriptome of mock-infected *Slpi^+/+^* (M-*wt*, n=4) and *Slpi^-/-^* (M-*ko*, n=3) and their UTI89-infected *Slpi^+/+^*(I-*wt*, n=14) and *Slpi^-/-^* (I-*ko*, n=11) counterparts. **B)** Volcano plot showing genes significantly upregulated in infected *Slpi^+/+^* (I-*wt*, blue) and infected *Slpi^-/-^* mice (I-*ko*, green). **C)** Bubble volcano plot showing gene set enrichment analysis of GO pathways between I-*wt* and I-*ko* mice. Differentially expressed child pathways are colored using the following categories: immune regulation (blue), DNA replication and repair (yellow), wound healing (orange), small molecule biosynthesis (pink), muscle growth (purple), neuronal regulation (green), or other (gray). GO pathways listed at the bottom are representative parent pathways for each color group. Pathways size is shown as bubble size (inset key). **D)** Enrichment plots (left) and count heatmaps (right) of inset labeled pathways from (**C**). Log_2_ fold change between I-*wt* and I-*ko* are shown in blue to green on the left, while normalized read counts for each group are shown as two columns to the right in purple. Significantly enriched genes as determined by DESeq2 are boxed. Five leading edge genes of each pathway that are either significantly enriched (boxed) and/or show the greatest enrichment are shown. Significantly enriched genes shown here are also labeled in (**B**). Note that gene *Lep* is listed twice as part of two different pathways. Dotted lines in all plots represent p value cutoff of 0.05.

We first contrasted infected I*-wt* and I*-ko* mice to their respective mock-infected control groups (M-*wt*, M-*ko*) to identify pathways significantly regulated by UTI. As expected, we found that infection triggered significant differential expression of 1878 genes in I-*wt* compared to M-*wt,* many of which are genes involved in the inflammatory response (Figure S3A; see also Table S1A). In *Slpi^-/-^*mice, only 257 genes were differentially expressed between I*-ko* and M*-ko* mice, of which 212 were shared with infected *Slpi^+/+^* mice (Table S1B). Interestingly, despite these differences in gene-level regulation, 71% of the pathways significantly regulated by infection were shared between *Slpi^+/+^* (I*-wt* vs. M*-wt*) and *Slpi^-/-^* (I*-ko* vs. M*-ko*) mice, suggesting that *Slpi^-/-^* mice respond to a UTI similarly to *Slpi^+/+^* (Figure S3A; see also Table S2A and S2B). To understand these contrasting findings, we examined pathways expected to be upregulated during a UTI that were also shared between infected *Slpi^+/+^* (I*-wt* vs. M*-wt*) and *Slpi^-/-^*(I*-ko* vs. M*-ko*) mice (Figure S3B). While the enrichment scores were similar between *Slpi^+/+^* (I*-wt* vs. M*-wt*) and *Slpi^-/-^* (I*-ko* vs. M*-ko*) groups, we found that many of the genes within these pathways were expressed more highly and these genes were more likely to be significant in *Slpi^+/+^* (I*-wt* vs. M*-wt*) mice (Figure S3C). These data show that while *Slpi^-/-^* mice respond to infection similarly to *Slpi^+/+^* mice at the pathway level, they significantly upregulated fewer individual inflammatory genes and appeared to have a muted inflammatory response at 1 dpi compared to wild-type controls.

To identify genes regulated by the presence of SLPI, we first compared mock-infected *Slpi^-/-^* and *Slpi^+/+^* mice (M-*ko* vs. M-*wt*) which demonstrated few significant differences at both the gene and pathway levels implying that in the absence of infection the transcriptional profiles of the bladder are very similar (Table S1C and S2C). We next contrasted I-*ko* to I-*wt* mice to investigate the impact of SLPI on the transcriptional response to UTI. We found that there were 42 differentially expressed genes, most of which were only modestly altered in expression (log2 fold change < 1; Figure 4B and S3D; see also Table S1D). We additionally verified by qRT-PCR that several inflammatory genes known to be important in UTI were not different between *Slpi^+/+^* and *Slpi^-/-^*mice at multiple time points (Figure S4A-C). However, pathway level analysis demonstrated 553 differentially expressed pathways, of which 508 were enriched in I-*wt* mice (Figure 4C; see also Table S2D). Most of the pathways differentially enriched in I-*wt* mice are involved in immune regulation, DNA replication and repair, small molecule biosynthesis, and wound healing (Figure 4C). These pathways encompass many of the genes found to be differentially expressed between I-*ko* to I-*wt* mice (Figure 4D). Together our findings suggest that the increased urine titers of UTI89 observed in *Slpi^-/-^* mice could be the result of a dysregulated immune response and delayed epithelial healing.

### *Slpi^-/-^* mice experience prolonged bladder inflammation after UTI

We next asked whether the absence of SLPI was associated with delayed resolution of bladder inflammation following UTI. We assessed this by collecting urines and bladders from *Slpi^+/+^* and *Slpi^-/-^* mice over a period of 7 hours to 7 days. H&E stained bladder sections were scored for inflammation severity on a scale of 0 to 5, where 0 was considered normal and 5 indicated severe damage to the uroepithelium (57). We found that *Slpi^-/-^* mice had significantly higher bladder inflammation scores at 2 and 7 dpi infection, although whole bladder weights did not differ at 3 hpi (Figure 5A and 5B; Figure S4D). Overall, there was a positive correlation between inflammation score and urine UPEC titers as expected for *Slpi^+/+^* and *Slpi^-/-^* mice (Figure 5C). Finally, we measured NE levels in the urine of mice infected with UPEC. We found NE in urine from infected *Slpi^-/-^* mice was significantly higher than infected *Slpi^+/+^* mice prior to infection, and at both early (3 hpi) and late timepoints (1 and 3 dpi) (Figure 5D), suggesting that SLPI may reduce the secretion and/or persistence of NE within the urinary tract even before infection.

**Figure 5:**
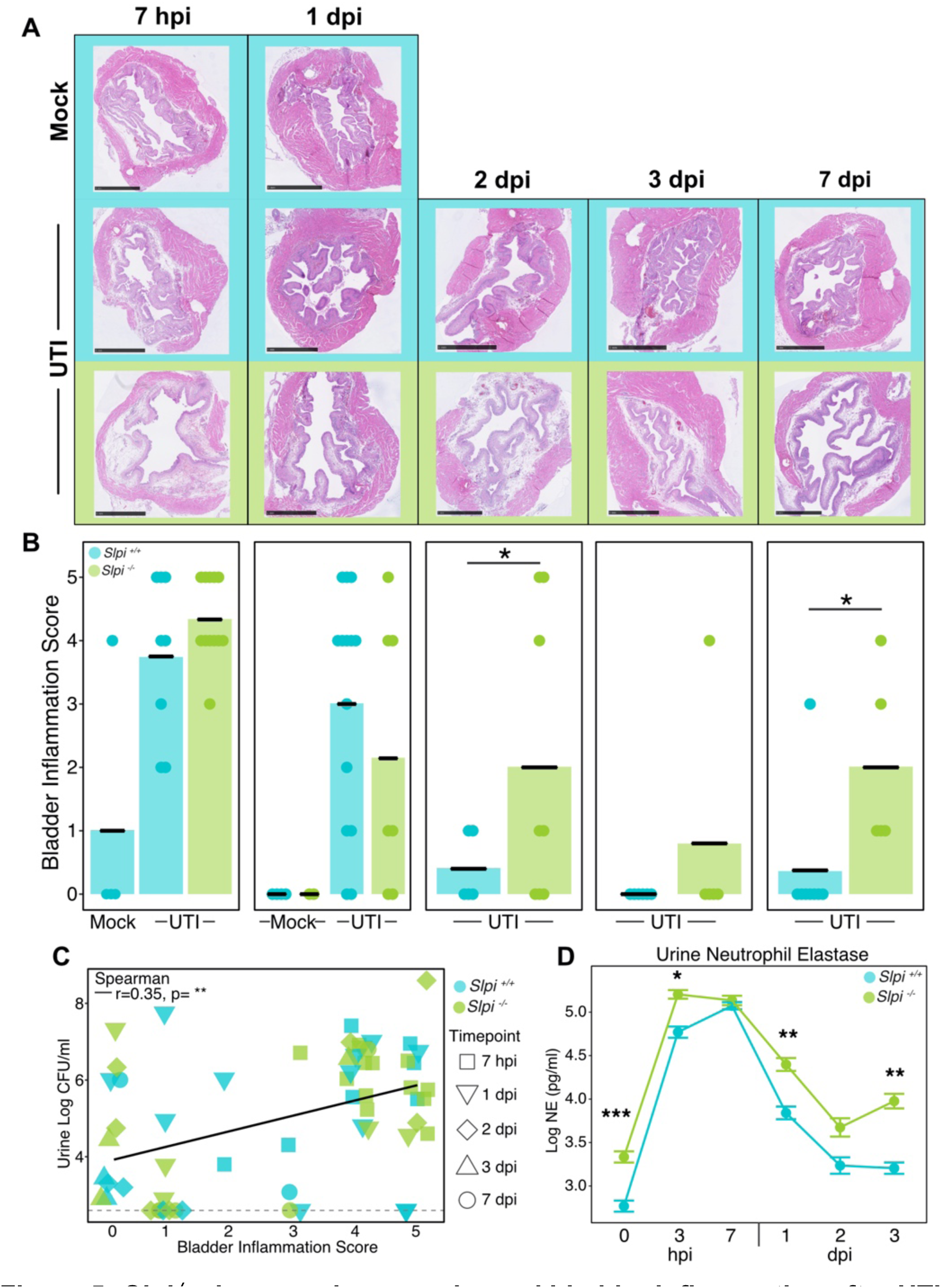
*Slpi^-/-^* mice experience prolonged bladder inflammation after UTI. **A)** Representative Hematoxylin and Eosin staining of bladders from mock-infected *Slpi^+/+^* (blue, top row), UTI89-infected *Slpi^+/+^* (blue, second row), and infected *Slpi^-/-^* mice (green, last row) at 7 hpi, 1, 2, 3, and 7 dpi. Images were taken at 20X magnification with 1 mm scalebars. **B)** Bladder inflammation scores (n=4-14) mice per group, combined from 5 experiments (one-sided Student’s t-test). **C)** Correlation of bladder inflammation score to urine log CFU/ml for all timepoints (n=2-12 per genotype per timepoint combined from 5 experiments; Spearman Rank). **D)** Log transformed neutrophil elastase (NE) levels in *Slpi^+/+^* (blue) and *Slpi^-/-^* mice (green) following infection. This data is combined from 6 experiments with 10-30 mice per genotype per timepoint (Student’s t-test).

### Levels of urine SLPI are changed in women with bacteriuria

Our mouse studies suggest that SLPI is increased in urine in response to UPEC-caused UTI, but the behavior of SLPI in the human urinary tract during UTI is not currently known. We addressed this gap in knowledge by obtaining urine and clinical information from female subjects 18-49 years old that had urine samples submitted to the clinical microbiology lab at Barnes-Jewish Hospital in St. Louis, Missouri. After collection, SLPI and total protein were quantified and correlated to the available clinical data.

We first excluded samples collected from pregnant individuals as well as samples from inpatient facilities to avoid potential cofounding medical conditions (see Methods). We also excluded samples that were contaminated or collected by urinary catheters. Of the remaining samples, we then removed samples from subjects with comorbidities that could influence urine SLPI levels including COVID-19, asthma, cardiovascular disease, cancer, diabetes, autoimmune disease, chronic kidney disease, and urinary anatomic abnormalities (including nephrolithiasis). The remaining 16 samples were then divided into three groups: 1) the first group had cultures negative for a uropathogen, no current UTI symptoms, and no reported history of a recent UTI within the last 6 months or rUTI (>2 infections in last month and/or >4 infections in any year prior); 2) the second group had cultures positive for a uropathogen (defined here as >10^5^ CFU/ml) and no recent history of UTI or rUTI; and 3) the third group had cultures positive for a uropathogen and a recent history of UTI or rUTI (Figure 6A). A one-way ANOVA showed these groups did not differ in age (p=0.9) or BMI (p=0.5) (Table S3). Comparison of these three groups showed that individuals with a culture positive for a uropathogen without a history of rUTI trended toward greater amounts of urine SLPI when compared to subjects with a negative culture (Figure 6B, p=0.06). However, women with a history of recent or rUTI with a culture positive for a uropathogen did not demonstrate any elevation in their urine SLPI levels compared to women without a cultured uropathogen. They also had significantly less urine SLPI when compared to uropathogen positive women without a history of recent or rUTI. Together, these results implicate SLPI in the response to uropathogen exposure but imply that its levels may be modulated by repeated exposure. Similarly, the uropathogen positive group trended higher in total urine protein compared to subjects with negative cultures, but this group was not significantly different from patients with a history of recent or rUTI (Figure 6C). This suggests that SLPI may be upregulated along with other proteins in response to a bacterial insult. We also examined available urinalysis results for these patients (Figure S5A) which includes biomarkers often used for diagnosis of a UTI including urine nitrite, leukocyte esterase (LE) and white blood cell count. However, we did not find any significant correspondence between any of these biomarkers and SLPI (Figure S5B-D).

**Figure 6:**
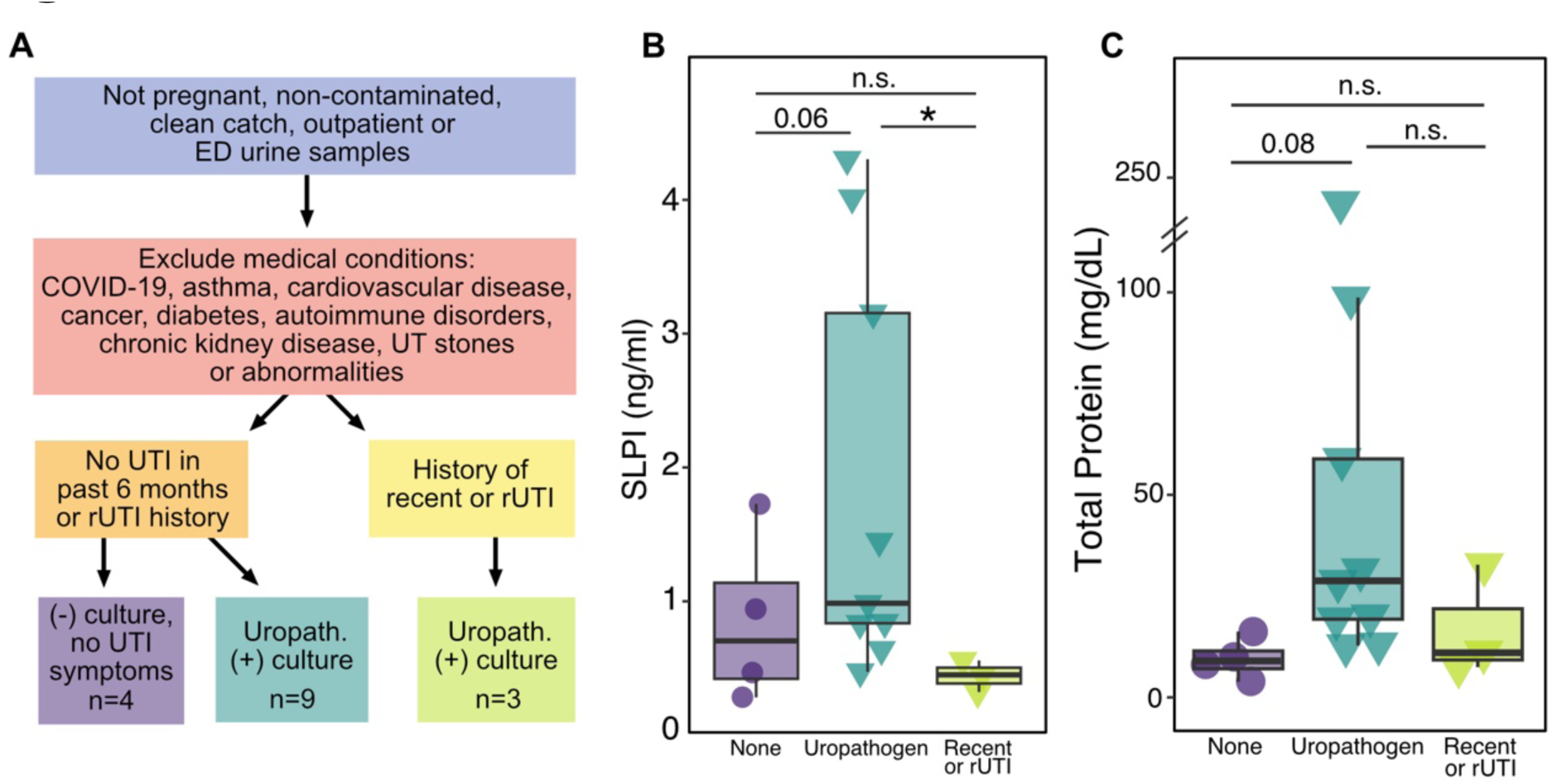
Levels of urine SLPI are changed in women with bacteriuria. **A)** Flowchart describing the filtering process for clinical study patient samples. **B)** SLPI protein measured in urine samples (one-sided Student’s t-test). **C)** Total protein in urine corresponding to samples in (B) (Student’s t-test). Shapes represent urine culture status: negative culture with no UTI symptoms (circles), culture positive for uropathogen (triangle).

## Discussion

SLPI is known to provide several protective effects to the host and has been directly demonstrated to aid in wound healing (28) and limit deleterious inflammation (58–60). While changes in SLPI expression have been associated with infections at mucosal sites, the relevance of SLPI during experimental infection is largely unexplored. Here, we use a well-characterized animal model of UTI (61) to show that SLPI is rapidly increased in the urine in response to UPEC challenge. Using a previously described mouse deficient in SLPI (*Slpi^-/-^*) (28), we show that mice lacking SLPI are more susceptible to severe UTI, as evidenced by dysregulation of the inflammatory response, increases in urine bacterial titers, as well as more severe and prolonged bladder inflammation. We also find that *Slpi^-/-^* mice have increased urine NE at baseline and during infection which likely contributes to the delayed histological resolution of infection. Our findings define SLPI as an important innate host factor that helps to repel bacterial pathogens within the urogenital tract, demonstrate a causal role for SLPI in resolving mucosal inflammation, and imply that SLPI may directly protect from infection at other mucosal sites.

SLPI is known to have immunomodulatory properties, including the ability to regulate production of NFkB-derived inflammatory cytokines (49,51,62) and modulate neutrophil function (63,64). Our observation that *Slpi^-/-^* mice demonstrate evidence of immune dysregulation at multiple stages of UPEC cystitis support an immune regulatory role for SLPI in the urinary tract. During the early stages of cystitis (7 hpi), we measured increased urine neutrophils in *Slpi^-/-^* mice but did not observe reduced bacterial titers at this same timepoint. At 1-2 dpi, *Slpi^-/-^* mice had reduced expression of inflammatory pathways compared to their *Slpi^+/+^* controls which was accompanied by increased urine titers of UPEC. Finally, during the resolution of infection, *Slpi^-/-^*mice showed delayed recovery of bladder epithelium following infection. Together, these findings suggest that in the absence of SLPI, there is a dysregulated immune response that peaks earlier than in *Slpi^+/+^* mice and also fails to effectively resolve bladder inflammation. These findings are in line with previous publications showing that controlling UPEC in the bladder requires precise regulation of neutrophil recruitment to ensure adequate clearance of bacteria while avoiding excessive inflammation that can lead to mucosal damage and chronic infection (24). Better understanding of the multipurposed role for SLPI in regulating neutrophil function, immune signaling, and epithelial recovery will require future experiments to untangle the tissue-specific roles of SLPI in the bladder.

Additionally, our findings suggest that the immunomodulatory and anti-serine protease activities of SLPI are not mutually exclusive. One of the major biological functions of SLPI is to inhibit the activity of NE, a serine protease that promotes inflammation by degrading extracellular matrix proteins especially elastin (43,44), causing increased permeability and excessive tissue damage (43,45–47,65). During experimental UTI, SLPI-deficient mice had increased NE (but not neutrophils, Figure 3B) in their urine even before infection and long after peak neutrophil infiltration in the urine, implying that SLPI limits NE abundance during homeostasis and infection in the urinary tract. This is potentially important because *Slpi^-/-^* mice had histologically more severe and prolonged inflammation in our experiments, which could be explained by an overabundance of NE found in the bladder lumen. Together, these findings imply that SLPI, similar to the serpin antiprotease family (24), helps to resolve inflammation in the bladder following infection by limiting the mucosal damaging protease activity of NE (64) in the urinary tract.

While multiple cell types are known to express SLPI, epithelial cells are primarily responsible for producing and secreting SLPI at mucosal surfaces (32,66,67). However, many immune cell types including macrophages (68), neutrophils (63,69), eosinophils (70), T (71) and B cells (72), express SLPI where it may play a direct role in regulating immune cell function (41). In the urinary tract, SLPI likely originates from multiple cell types, including the bladder epithelium, and is also known to be directly filtered and secreted into the urine from the circulation through the kidneys (31,73–75). Our study shows that SLPI is localized to bladder epithelial cells, suggesting urine SLPI is arising, at least in part, from the bladder epithelium during infection. Increases in SLPI seen in the urine could come through enhanced secretion, exfoliation and lysis of infected cells, or a combination of both. Interestingly, rises in SLPI protein found in urine and cell culture supernatants following infection is not accompanied by corresponding increases in *Slpi* RNA transcripts. These findings raise the possibility that bladder cells contain a reservoir of SLPI that is rapidly released following pathogen challenge rather than transcriptionally regulated during the early stages of infection. Additionally, we cannot exclude the possibility that SLPI may also originate from recruited neutrophils and immune cells in our mouse model, though we note that neutrophil abundance in the urine typically subsides before SLPI protein levels return to baseline (76). Future experiments with a tissue-specific deletion of SLPI will further elucidate the cellular origins of SLPI in the urinary tract during infection.

In addition to its antiprotease and immunomodulatory activities, SLPI has been reported to inhibit the growth of *E. coli* (38), *S. aureus* (38), Group A *Streptococcus* (77), and *P. aeruginosa* (55) in culture, and direct antimicrobial activity against UPEC would certainly explain our observation of increased urine titers in *Slpi^-/-^* mice during UTI. However, we did not detect bactericidal activity of commercially-obtained recombinant SLPI (rSLPI) against the prototypical UPEC strain, UTI89, or a lab adapted strain of *E. coli* in our *in vitro* studies. While we cannot exclude the possibility that SLPI inhibits UPEC growth under different *in vitro* conditions or only *in vivo*, cathelicidin (LL-37), another antimicrobial peptide found in the urinary tract (29), demonstrated high potency in the same assay. Our results are consistent with previous findings of *P. aeruginosa* infection of the airway (78) and suggest that the immunomodulatory functions of SLPI mediate its protective effects in our model of UTI.

We also investigated the role of SLPI in the human urinary tract by analyzing SLPI abundance in the urine of a cross-sectional group of women. When we focused on a subset of non-pregnant women without significant comorbidities (including prior UTI) that had negative urine bacterial growth or were positive for a uropathogen, we found SLPI tended to be increased in uropathogen-positive urine cultures. This finding supports our conclusions from our mouse experiments that SLPI initially increases rapidly in the urine in response to uropathogen invasion. However, we also noticed that women with a history of rUTI or recent UTI did not increase urine SLPI in the presence of a uropathogen. While this may appear to contradict our observations from our animal model, SLPI at other mucosal sites, like the lung, have been noted to decline in conditions of chronic inflammation and other antiproteases have been noted to have reduced expression in the bladder following rUTI (79,80). Additionally, we note that decreases in urine SLPI have been described by other investigators in patients experiencing recurrent or frequent UTIs (75). These prior studies combined with our current findings suggest that SLPI may be rapidly released from bladders that are naive to infections, but that SLPI levels may decline with repeated or prolonged inflammation. This could be potentially explained by a reservoir of SLPI residing in the bladder epithelium that is rapidly released during an acute infection but is depleted with recurrent stimulation. Our immunohistochemistry (Figure 2) supports this idea, showing that staining for SLPI within the bladder epithelium appears to be diminished in mice after experiencing UTI despite high levels of SLPI in the urine. Future studies dissecting the temporal relationship of urine SLPI to UTI, particularly in the context of recurrent UTI, will be necessary to untangle the relationship between SLPI, UTI, and chronic inflammation.

There are several limitations to this study that should be considered. First, while the *Slpi^-/-^* mice used in our study were backcrossed 10+ generations, we cannot exclude the possibility that passenger mutations (81) affecting genes in regions flanking *Slpi* (e.g. *Serinc3*, *Pkig*) indirectly influence our phenotypes. Reassuringly, in a reanalysis of previously published data, we found that many of the genes that were differentially expressed during UTI were also regulated by SLPI when it was over expressed in mouse preosteoblast cells (82) (Figure S6), suggesting that our findings are independent of potential confounding passenger mutations. Second, in our clinical study, we do not have data about whether patients were treated for UTI, so we are uncertain if subjects with a cultured uropathogen were ultimately diagnosed with a UTI. Third, the subject population tended to have multiple comorbidities in addition to UTI and samples were collected during periods of high community transmission of COVID-19 infection. Additionally, approximately half of our samples were made up of routine urine screenings for pregnant women which were excluded from our analyses of SLPI and uropathogens (see Methods). As a result, the study population on which we could perform analyses was significantly reduced.

In summary, the early innate immune response to bacterial pathogens can set the stage for progression, persistence, or resolution of infection. Our findings show that SLPI restrains pathogen levels while simultaneously limiting inflammation, helping to achieve a delicate balance between the immune response and protection of the bladder tissues. We suggest that further understanding of the diverse roles SLPI plays in the urinary tract could improve our understanding and treatment of UTIs.

## Materials and Methods

### Mice

All animal procedures were reviewed by the Washington University Institutional Animal Care and Use Committee (Protocol#: 21-0394). Female WT C57BL/6J mice were purchased from Jackson Laboratories (CAS: 00064, Bar Harbor, ME). Cryo-recovered 129;BL/6 SLPI^tm1Smw^/J mice (CAS: 010926, Jackson Laboratories, Bar Harbor, ME) were backcrossed to WT C57BL/6J mice for at least 10 generations before being used in these experiments. Female *Slpi^+/+^*and *Slpi^-/-^* mice were generated through homozygous breeding.

### Mouse Urinary Tract Infection

A kanamycin-resistant derivative of a human cystitis isolate, UTI89*att_HK202_::KanR* (called UTI89 throughout this manuscript) and used to infect mouse bladders (52). This strain was a generous gift from Fimbrion Therapeutics, Inc. (St. Louis, MO) and was generated by inserting a kanamycin resistance cassette into the HK022 phage bacterial attachment site of the human cystitis UPEC isolate UTI89 (12) using the lambda red recombinase system, as previously described (83,84). The insertion cassette was amplified from pKD4 using primers HK-FT-F and HK-FT-R, producing a linear PCR product with ends that are homologous to HK site sequences (Table S4A). The linear PCR product was then electroporated into UTI89 containing a temperature sensitive helper plasmid pKM208 and plated on agar containing kanamycin. Recombinants were confirmed by PCR using flanking primers HK-test-Left and HK-test-Right to confirm insertion at the HK site (Table S4A).

Bacterial cultures were grown under type 1 pilus inducing conditions. Briefly, 20 ml of Luria-Bertani broth (LB) was seeded from a glycerol stock and grown statically at 37°C overnight, then subcultured 1:1000 in fresh LB, and grown statically at 37°C for an additional 18-24 hours. Cultures were centrifuged (1,750g for 15 min) and resuspended in sterile 1X PBS. Inoculum was diluted to a final concentration of 1-2×10^9^ colony forming units (CFU)/ml (1:10 OD600=0.22-0.24) and 50 μl (∼0.8-1×10^8^ CFU) was used to inoculate the bladders of 6-8 week old female mice. Transurethral catheterization was performed while mice were anesthetized using inhaled 2.5% isoflurane. Mock-infected mice were inoculated with sterile PBS in the same manner.

### Bacterial Enumeration in Mouse Tissues

Following infection, mice were euthanized then bladder and kidneys were aseptically harvested into pre-weighed tubes containing 1 ml of sterile PBS. In some experiments, bladders were cut in half longitudinally with one section being saved for other assays. Final weights were recorded before homogenizing tissues. Samples (including urine) were serially diluted 1:10 in sterile PBS and 5 μl of each dilution was spotted onto LB agar supplemented with 50 μg/ml kanamycin five times. Colony counts were averaged for the highest dilution containing colonies in 4 or more replicates and CFU/ml was calculated. This was normalized to total tissue weight to obtain CFU/g.

### *Ex vivo* Gentamicin Protection Assay

An *ex vivo* gentamicin protection assay (54) was used to determine the amount of attached and intracellular UPEC with some modifications. Briefly, after 3 hours of infection, half bladders were removed from mice and washed 3 times with 500 μl sterile PBS with the second wash used to quantify luminal UPEC. Bladders were then placed in PBS with gentamicin (100 ug/ml) and incubated at 37°C for 90 minutes. After washing with 1 ml of PBS, bladders were homogenized in 0.1% triton/PBS, diluted, and plated on LB/Kan50 agar plates to determine the amount of intracellular UPEC.

### Neutrophil Quantification by Cytospin

Urine was diluted 1:10 and 80 μl were spun onto poly-L-lysine-coated glass slides (750g, 10 min). Slides were air-dried, fixed for at least 1 hour, and stained using the Shandon KwikDiff staining kit (CAS: 99-907-01, Thermofisher). Coverslips were mounted using Cytoseal XYZ (CAS: 22-050-262, Thermofisher) and allowed to dry. Investigator was blinded to slides before neutrophils were counted across 3 high power fields (HPF; 40X magnification) and then averaged for each sample.

### Protein Quantification by ELISA

Urine and 5637 cell culture supernatant samples were thawed and diluted 1:10 (uninfected and mock-infected) or 1:200-1:500 (infected) in sterile PBS + 1% BSA. SLPI and neutrophil elastase (NE) were measured using the Mouse SLPI DuoSet ELISA (CAS: DY1735-05, R&D Systems) and Mouse Neutrophil Elastase (ELA2) Duoset ELISA (CAS: DY4517-05, R&D Systems) respectively, both according to manufacturer’s instructions.

### Bacterial killing assay

UTI89 and DH5ɑ were each shaking at 37°C in 5 ml of LB broth and then subcultured into a fresh 5 ml of LB. This culture was grown to log phase (∼2 hours) before being centrifuged (1,750g, 15 min), washed once with sterile 1X PBS, and resuspended to 1:10 OD600=0.15-0.2. The culture was then diluted 1:50 in 1X PBS + 1% LB broth and incubated statically at 37°C with recombinant human SLPI protein (rSLPI) (CAS: 1274-PI-100, R&D Systems) at these final concentrations: 20μM, 10μM, 1μM. A 1μM concentration of LL-37 (CAS: tlrl-l37, Invivogen), the active portion of cathelicidin, was used as a positive control for bactericidal activity. Cultures were sampled after 2 hours of incubation and titered as previously described.

### Bladder Histological Scoring

Half bladders were stored in 4% paraformaldehyde/water at 4°C overnight. The following day, samples were washed 3 times with 70% ethanol and then stored at 4°C until further processing. Samples were pinned to silicone plates, embedded in 1% agar, and stored in 70% ethanol until paraffin embedding and hematoxylin and eosin (H&E) staining was performed. Slides were then imaged using a Hamamatsu Nanozoomer at 20X power. An investigator, who was blinded to timepoints and experimental groups, assigned a score of 0-5 to each bladder using a grading system adapted to the robust UTI response seen in C57BL/6 mice (19). Scores used in this manuscript are defined as the following: 0, Normal; 1, Focal neutrophil infiltration; 2, Diffuse neutrophil infiltration; 3, Diffuse neutrophil infiltration and edema for early acute cystitis; 4, All of 3 plus neutrophils in muscle tissue; 5, Erosion of bladder epithelium.

### Bladder Immunofluorescent Staining

Bladders were harvested and processed for histology as described above. Unstained paraffin-embedded sections underwent paraffin removal using 100% xylenes followed by 100% ethanol to remove excess xylene. Subsequent 5 minute incubations in decreasing concentrations of ethanol (95%, 90%, 70%, and 50%) were used to rehydrate the tissue followed by water to remove residual ethanol. For antigen retrieval, a 10 mM sodium citrate + 0.05% Tween20, pH 6.0 solution was used in which slides were microwave-boiled 3 times then left in the warm buffer for 30 minutes before being washed 3X in water. Samples were permeabilized for 10 minutes with 0.1% Triton-×100 in PBS, washed 3X with PBS, incubated 30 minutes with 0.1% TrueBlack Plus Lipofuscin Autofluorescence Quencher (CAS: 23014, Biotium) then washed 3X with PBS. Slides blocked for 30 minutes with 2% BSA+2% donkey serum in PBS then washed 3X with PBS. Then, samples were incubated in 5 μg/ml polyclonal rabbit anti-mouse SLPI antibody (CAS: NBP1-76803, Novus Biologicals) or rabbit polyclonal IgG isotype control (CAS: NBP2-24891, Novus Biologicals) diluted in PBS+1% normal donkey serum (CAS: 017-000-121, Jackson ImmunoResearch) overnight at 4°C. The following day, samples were washed 3X in PBS before incubating 2 hours at room temperature in 2 μg/ml donkey anti-rabbit IgG conjugated to AF594 (CAS: A21207, Thermofisher). After 3 washes in PBS, a 5 μg/ml of Hoechst 33342 stain (CAS: H21492, Invitrogen) in PBS was added before a final 3 washes with PBS and coverslip mounting using ProLong Diamond Antifade Mountant (CAS: P36961, Thermofisher). Slides were imaged at 5X and 20X using a Zeiss Axio Observer Microscope and fluorescence display settings were determined using isotype and infected *Slpi^-/-^* mouse controls.

### Bladder RNA isolation

Half bladder samples stored in 500 μl of RNAlater (CAS: AM7021, Thermofisher) incubated overnight at 4°C. The following day, RNAlater was removed and samples transferred to -80°C until further processing. To isolate RNA, bladders were thawed, transferred to 1 ml of Trizol and homogenized. A 0.3 ml portion of this homogenate was used to extract and purify RNA using the RNeasy Mini Kit (CAS: 74104, Qiagen) following manufacturer’s instructions, then eluted in 25 μl of RNAse-free water.

### Quantitative PCR on bladder homogenate

After extraction, RNA quality was checked by gel electrophoresis and quantified using Ribogreen (CAS: R11490, Invitrogen). High-Capacity cDNA Reverse Transcription Kit (CAS: 4387406, Applied Biosystems) was used to reverse transcribe 100 ng of RNA to cDNA according to manufacturer’s instructions. cDNA was diluted 1:3 or 1:4 before mixing with Power SYBR Green PCR Master Mix (CAS: 4368702, Applied Biosystems) and primer pairs (Table S4B). Results were analyzed using the ddCT method(85).

### Bulk RNA sequencing on whole bladder

Following extraction, RNA concentration was determined using Qubit RNA Quantification Assay, High Sensitivity (CAS: Q32852, Invitrogen) before assessing RNA quality by BioAnalyzer (Agilent) to ensure all samples had a minimum RNA integrity number (RIN) of 8. Stranded, poly-A enriched libraries were created using the NEBNext Poly(A) mRNA Magnetic Isolation Module (CAS: E7490S, NEB) followed by the NEBNext Ultra II Directional RNA Library Prep Kit (CAS: E7760S, NEB). Completed libraries were then sequenced to an average depth of approximately 20M reads per sample on a partial lane of the NovaSeq6000 S4 XP flow cell using 2×150 paired-end reads with 10-base dual indexes (CAS: E6440S, NEB). After demultiplexing these samples, reads were mapped to the Ensembl release 109 cDNA database using Salmon with default parameters (86). Differentially expressed genes were identified using *DESeq2* (v1.34.0) (87) and then mapped to entrez ID using *biomaRt* (v2.50.3) (88). Functional pathway analysis was performed using *fgsea* (v1.20.0) (89) and mouse biological process GO database to determine significantly altered pathways between groups.

### Bladder Cell Culture

Human bladder epithelial cells (5637) were maintained in RPMI 1640 (CAS: 11875085, Gibco) with 10% FBS (CAS: 16000-014, Gibco) in a humidified chamber set at 37°C with 5% CO_2_. Bacterial cultures for these experiments were grown for type 1 pilus expression as described above. The day prior to infection, 24-well plates were seeded with 2×10^5^ cells/well in 0.5 ml and incubated in the chamber overnight. The following day, 95-98% cell confluency was confirmed and media was replaced 3-4 hour before adding 50 μl of bacteria (4-7×10^6^ CFU, MOI 18-30) or PBS, then plates were gently centrifuged to settle bacteria onto cells before incubating at 37°C with 5% CO_2_. A 50 μl sample of supernatant was removed at 0, 0.5, and 1 hour of infection with remaining supernatant collected at 2 hours. At this time, cells were resuspended in Trizol reagent and frozen at -20°C until RNA was extracted using the Macherey-Nagel Nucleospin RNA XS Kit (CAS: 740902.50, Takara Bio) according to manufacturer’s instructions. A 250 ng input of RNA was used to make cDNA which was subsequently diluted 1:5 in nuclease-free water before being used in qPCR as described above.

### SLPI-UTI Clinical Study

This study was approved by the Institutional Review Board of Washington University in St. Louis (Protocol#: 202107050) which was granted a waiver of consent. It was designed to investigate the contribution of SLPI in urinary tract infection. Leftover urine used in this study was sourced from original samples submitted to a clinical microbiology lab for routine culturing and/or urinalysis testing and consent was waived. Inclusion criteria required St. Louis Barnes Jewish or Children’s Hospital patients to be female and 18-49 years old. Samples were collected between August 2021 and February 2022. Urine was aliquoted in 1 ml portions and stored at -80°C until further processing. SLPI was measured urine samples diluted 1:20-1:200 in sterile PBS + 1% BSA using the Human SLPI DuoSet ELISA kit (CAS: DY1274-05, R&D Systems) according to the manufacturer’s instructions. Total protein in the urine was measured using Quantichrom Total Protein Assay Kit (CAS: QTPR-100, BioAssay Systems) according to the manufacturer’s instructions with samples detected at a 1:4-1:20 dilution. A total of 202 patient samples were collected of which total protein and SLPI were measured in 197 samples. We defined a recent UTI as a documented infection within the last 6 months. Recurrent UTI (rUTI) was defined as more than 2 infections within the month before sampling and/or more than 4 infections within the last year.

We noted that pregnant women, who constituted 48.7% of the available urine samples, had significantly increased levels of SLPI in their urine compared to non-pregnant women (Table S2, Figure S7A). Given these differences, we excluded pregnant women from our analysis of uropathogens and SLPI.

### Statistics

Statistical analysis was performed using R Version 4.1.2 (R Development Core Team, 2021). Data are presented as mean with error bars denoting SEM, box and whisker plots, or scatterplots. Log_10_ transformed data prior to mean and SEM calculations is indicated in figure axis. Unless otherwise stated, statistical significance was conducted using unpaired Wilcoxon or Student’s t-test. Boxplots show IQR and whiskers display 1.5*IQR. Time series ANOVA was performed using *Amelia* (v1.8.0) package in R to impute missing data. Associations were determined using Spearman’s Rank. In all figures, the following symbols were used to designate significance: n.s. = not significant, * p<0.05, ** p<0.01, *** p<0.001, and **** p<0.0001.

### Data Availability

RNAseq data has been deposited at the European Nucleotide Archive (https://www.ebi.ac.uk/ena/browser/home) and is publicly available as of the date of publication under project accession number PRJEB66138.

## Acknowledgements

We would like to thank Jennifer Hazen and Benjamin Olson for their aid with the mouse model of UTI. We would also like to acknowledge Philip Ahern for critical feedback about our animal models and Tarisa Mantia for her assistance with setting up the clinical study. Further, we thank Rachel Kinsella for her guidance on immunofluorescence staining, the Alafi Neuorimaging Laboratory the Hope Center for Neurological Disorders, and NIH shared Instrumentation for use of the Hamamatsu Nanozoomer microscope (S10 RR0227552) to Washington University. Andrew L. Kau is supported in part by the Doris Duke Foundation (2019083). The National Institutes of Health provided support to Nicole M. Gilbert (NIDDK K01 DK110225), Jesús Santiago Borges (T32 GM007067), Thomas J. Hannan (U01AI095542), as well as the Hultgren lab (R01DK132327, U01AI095542), and the Kau lab (R01AI165915).

## Author Contributions

A.L.R. and A.L.K. conceptualized the work. A.L.R., M.A.W., C.D.B. and A.L.K. planned the clinical study and D.H.V. performed the chart review. A.L.R., M.A.L., D.H.V., N.M.G., C.P.T., J.S.B., S.J.H., and A.L.K. contributed to the design and conduct of experiments. M.A.L. and C.P.T. backcrossed and maintained all transgenic mice. T.J.H. performed the blinded scoring of bladders. A.L.R. and A.L.K. drafted the manuscript. All authors interpreted the data and contributed to revising the manuscript.

## References

1. Griebling TL. Urologic diseases in America project: Trends in resource use for urinary tract infections in women. Journal of Urology [Internet]. 2005 [cited 2021 Apr 12];173(4):1281–7. Available from: https://pubmed.ncbi.nlm.nih.gov/15758783/

2. Flores-Mireles AL, Walker JN, Caparon M, Hultgren SJ. Urinary tract infections: Epidemiology, mechanisms of infection and treatment options [Internet]. Vol. 13, Nature Reviews Microbiology. Nature Publishing Group; 2015 [cited 2021 Apr 12]. p. 269–84. Available from: https://pubmed.ncbi.nlm.nih.gov/25853778/

3. Najar MS, Saldanha CL, Banday KA. Approach to urinary tract infections [Internet]. Vol. 19, Indian Journal of Nephrology. Indian J Nephrol; 2009 [cited 2021 Apr 13]. p. 129–39. Available from: https://pubmed.ncbi.nlm.nih.gov/20535247/

4. Kalinderi K, Delkos D, Kalinderis M, Athanasiadis A, Kalogiannidis I. Urinary tract infection during pregnancy: current concepts on a common multifaceted problem [Internet]. Vol. 38, Journal of Obstetrics and Gynaecology. Taylor and Francis Ltd; 2018 [cited 2021 Apr 13]. p. 448–53. Available from: https://pubmed.ncbi.nlm.nih.gov/29402148/

5. Scholes D. Risk factors for recurrent urinary tract infection in young women. Journal of Infectious Diseases [Internet]. 2000 [cited 2021 Apr 12];182(4):1177–82. Available from: https://pubmed.ncbi.nlm.nih.gov/10979915/

6. Foxman B. Urinary tract infection syndromes. Occurrence, recurrence, bacteriology, risk factors, and disease burden [Internet]. Vol. 28, Infectious Disease Clinics of North America. Infect Dis Clin North Am; 2014 [cited 2021 Apr 11]. p. 1–13. Available from: https://pubmed.ncbi.nlm.nih.gov/24484571/

7. Svanborg C, Godaly G. Bacterial virulence in urinary tract infection. Infect Dis Clin North Am [Internet]. 1997 [cited 2021 Apr 12];11(3):513–29. Available from: https://pubmed.ncbi.nlm.nih.gov/9378921/

8. Recurrent Urinary Tract Infections in Female Patients : An Overview of Management and Treatment [with Panel Discussion] Author (s): Thomas A . Stamey Source : Reviews of Infectious Diseases, Vol . 9, Supplement 2. Update and Advances in Intravenous. 2018;9.

9. Barber AE, Norton JP, Wiles TJ, Mulvey MA. Strengths and Limitations of Model Systems for the Study of Urinary Tract Infections and Related Pathologies. Microbiology and Molecular Biology Reviews [Internet]. 2016 Jun [cited 2021 Apr 12];80(2):351–67. Available from: https://pubmed.ncbi.nlm.nih.gov/26935136/

10. Mulvey MA, Lopez-Boado YS, Wilson CL, Roth R, Parks WC, Heuser J, et al. Induction and evasion of host defenses by type 1-piliated uropathogenic Escherichia coli. Science (1979). 1998;282(5393):1494–7.

11. Martinez JJ, Mulvey MA, Schilling JD, Pinkner JS, Hultgren SJ. Type 1 pilus-mediated bacterial invasion of bladder epithelial cells. EMBO Journal. 2000;19(12):2803–12.

12. Mulvey MA, Schilling JD, Hultgren SJ. Establishment of a persistent Escherichia coli reservoir during the acute phase of a bladder infection. Infect Immun. 2001;69(7):4572–9.

13. Justice SS, Hung C, Theriot JA, Fletcher DA, Anderson GG, Footer MJ, et al. Differentiation and developmental pathways of uropathogenic Escherichia coli in urinary tract pathogenesis. Proc Natl Acad Sci U S A [Internet]. 2004 Feb 3 [cited 2021 Apr 13];101(5):1333–8. Available from: https://pubmed.ncbi.nlm.nih.gov/14739341/

14. Anderson GG, Palermo JJ, Schilling JD, Roth R, Heuser J, Hultgren SJ. Intracellular bacterial biofilm-like pods in urinary tract infections. Science (1979) [Internet]. 2003 Jul 4 [cited 2021 Apr 13];301(5629):105–7. Available from: https://pubmed.ncbi.nlm.nih.gov/12843396/

15. Thumbikat P, Berry RE, Schaeffer AJ, Klumpp DJ. Differentiation-induced uroplakin III expression promotes urothelial cell death in response to uropathogenic E. coli. Microbes Infect. 2009;11(1):57–65.

16. Kuhn HW, Hreha TN, Hunstad DA. Immune defenses in the urinary tract. Vol. 44, Trends in Immunology. Elsevier Ltd; 2023. p. 701–11.

17. Ashkar AA, Mossman KL, Coombes BK, Gyles CL, Mackenzie R. FimH adhesin of type 1 fimbriae is a potent inducer of innate antimicrobial responses which requires TLR4 and type 1 interferon signalling. PLoS Pathog. 2008 Dec;4(12).

18. Ali ASM, Mowbray C, Lanz M, Stanton A, Bowen S, Varley CL, et al. Targeting Deficiencies in the TLR5 Mediated Vaginal Response to Treat Female Recurrent Urinary Tract Infection. Sci Rep. 2017 Dec 1;7(1).

19. Hannan TJ, Mysorekar IU, Hung CS, Isaacson-Schmid ML, Hultgren SJ. Early severe inflammatory responses to uropathogenic E. coli predispose to chronic and recurrent urinary tract infection. PLoS Pathog [Internet]. 2010 Aug [cited 2021 Apr 13];6(8):29–30. Available from: https://pubmed.ncbi.nlm.nih.gov/20811584/

20. Ingersoll MA, Kline KA, Nielsen H V., Hultgren SJ. G-CSF induction early in uropathogenic Escherichia coli infection of the urinary tract modulates host immunity. Cell Microbiol [Internet]. 2008 [cited 2021 Apr 13];10(12):2568–78. Available from: https://pubmed.ncbi.nlm.nih.gov/18754853/

21. Hayes BW, Abraham SN. Innate Immune Responses to Bladder Infection. Microbiol Spectr [Internet]. 2016 Dec 1 [cited 2021 Apr 13];4(6). Available from: https://pubmed.ncbi.nlm.nih.gov/28084200/

22. Shahin RD, Engberg I, Hagberg L, Eden CS. Neutrophil recruitment and bacterial clearance correlated with LPS responsiveness in local gram-negative infection . □ NEUTROPHIL RECRUITMENT AND BACTERIAL CLEARANCE CORRELATED WITH LPS RESPONSIVENESS IN LOCAL GRAM-NEGATIVE INFECTION ’. 1987;

23. Haraoka M, Hang L, Rn Frendéus B, Godaly G, Burdick M, Strieter R, et al. Neutrophil Recruitment and Resistance to Urinary Tract Infection [Internet]. Available from: https://academic.oup.com/jid/article/180/4/1220/843232

24. Hannan TJ, Roberts PL, Riehl TE, van der Post S, Binkley JM, Schwartz DJ, et al. Inhibition of cyclooxygenase-2 prevents chronic and recurrent cystitis. EBioMedicine. 2014;1(1):46–57.

25. Valore E V, Park CH, Quayle AJ, Wiles KR, Mccray PB, Ganz T. Human ␤ - Defensin-1 : An Antimicrobial Peptide of Urogenital Tissues. 1998;

26. Morrison G, Kilanowski F, Davidson D, Dorin J. Characterization of the Mouse Beta Defensin 1, Defb1, Mutant Mouse Model. 2002;70(6):3053–60.

27. Ali ASM, Townes CL, Hall J, Pickard RS. Maintaining a Sterile Urinary Tract: The Role of Antimicrobial Peptides [Internet]. Vol. 182, Journal of Urology. J Urol; 2009 [cited 2021 Apr 13]. p. 21–8. Available from: https://pubmed.ncbi.nlm.nih.gov/19447447/

28. Ashcroft GS, Lei K, Jin W, Longenecker G, Kulkarni AB, Greenwell-Wild T, et al. Secretory leukocyte protease inhibitor mediates non-redundant functions necessary for normal wound healing. Nat Med [Internet]. 2000 [cited 2021 Apr 13];6(10):1147–53. Available from: https://pubmed.ncbi.nlm.nih.gov/11017147/

29. Chromek M, Slamová Z, Bergman P, Kovács L, Podracká L, Ehrén I, et al. The antimicrobial peptide cathelicidin protects the urinary tract against invasive bacterial infection. Nat Med [Internet]. 2006 Jun [cited 2021 Apr 13];12(6):636–41. Available from: https://pubmed.ncbi.nlm.nih.gov/16751768/

30. Nielsen KL, Dynesen P, Larsen P, Jakobsen L, Andersen PS, Frimodt-møller N. Role of Urinary Cathelicidin LL-37 and Human ␤ -Defensin 1 in Uncomplicated Escherichia coli Urinary Tract Infections. 2014;82(4):1572–8.

31. Chaudhry R, Madden-Fuentes RJ, Ortiz TK, Balsara Z, Tang Y, Nseyo U, et al. Inflammatory response to Escherichia coli urinary tract infection in the neurogenic bladder of the spinal cord injured host. Journal of Urology [Internet]. 2014 [cited 2021 Apr 13];191(5):1454–61. Available from: https://pubmed.ncbi.nlm.nih.gov/24342147/

32. Kommunehospital A. GYNE Secretory leukocyte protease inhibitor in the cervical mucus and in the fetal membranes. 1995;59:95–101.

33. McKiernan PJ, McElvaney NG, Greene CM. SLPI and inflammatory lung disease in females. Biochem Soc Trans. 2011;39(5):1421–6.

34. Ma G, Greenwell-Wild T, Lei K, Jin W, Swisher J, Hardegen N, et al. Secretory leukocyte protease inhibitor binds to annexin II, a cofactor for macrophage HIV-1 infection. Journal of Experimental Medicine [Internet]. 2004 Nov 15 [cited 2021 Apr 13];200(10):1337–46. Available from: https://pubmed.ncbi.nlm.nih.gov/15545357/

35. Cooper MD, Roberts MH, Barauskas OL, Jarvis GA. Secretory Leukocyte Protease Inhibitor Binds to Neisseria gonorrhoeae Outer Membrane Opacity Protein and is Bactericidal. American Journal of Reproductive Immunology [Internet]. 2012 Aug [cited 2021 Apr 13];68(2):116–27. Available from: https://pubmed.ncbi.nlm.nih.gov/22537232/

36. Curvelo JADR, Barreto ALS, Portela MB, Alviano DS, Holandino C, Souto-Padrón T, et al. Effect of the secretory leucocyte proteinase inhibitor (SLPI) on Candida albicans biological processes: A therapeutic alternative? Arch Oral Biol [Internet]. 2014 [cited 2021 Apr 13];59(9):928–37. Available from: https://pubmed.ncbi.nlm.nih.gov/24907522/

37. Bingle CD, Vyakarnam A. Novel innate immune functions of the whey acidic protein family [Internet]. Vol. 29, Trends in Immunology. Trends Immunol; 2008 [cited 2021 Apr 13]. p. 444–53. Available from: https://pubmed.ncbi.nlm.nih.gov/18676177/

38. Hiemstra PS, Maassen RJ, Stolk J, Heinzel-Wieland R, Steffens GJ, Dijkman JH. Antibacterial activity of antileukoprotease. Infect Immun [Internet]. 1996 [cited 2021 Apr 13];64(11):4520–4. Available from: https://pubmed.ncbi.nlm.nih.gov/8890201/

39. Koizumi M, Fujino A, Fukushima K, Kamimura T, Takimoto-Kamimura M. Complex of human neutrophil elastase with 1/2SLPI. J Synchrotron Radiat [Internet]. 2008 Apr 18 [cited 2021 Apr 13];15(3):308–11. Available from: https://pubmed.ncbi.nlm.nih.gov/18421166/

40. Yang J, Zhu J, Sun D, Ding A. Suppression of macrophage responses to bacterial lipopolysaccharide (LPS) by secretory leukocyte protease inhibitor (SLPI) is independent of its anti-protease function. Biochim Biophys Acta Mol Cell Res [Internet]. 2005 Sep 30 [cited 2021 Apr 13];1745(3):310–7. Available from: https://pubmed.ncbi.nlm.nih.gov/16112212/

41. Majchrzak-Gorecka M, Majewski P, Grygier B, Murzyn K, Cichy J. Secretory leukocyte protease inhibitor (SLPI), a multifunctional protein in the host defense response [Internet]. Vol. 28, Cytokine and Growth Factor Reviews. Elsevier Ltd; 2016 [cited 2021 Apr 13]. p. 79–93. Available from: https://pubmed.ncbi.nlm.nih.gov/26718149/

42. Dewald B, Rindler Ludwig R, Bretz U, Baggiolini M. Subcellular localization and heterogeneity of neutral proteases in neutrophilic polymorphonuclear leukocytes. Journal of Experimental Medicine. 1975;141(4):709–23.

43. Döring G. The role of neutrophil elastase in chronic inflammation. Am J Respir Crit Care Med. 1994;150(6 II).

44. Morris SM, Stone PJ. Immunocytochemical study of the degradation of elastic fibers in a living extracellular matrix. Journal of Histochemistry and Cytochemistry. 1995;43(11):1145–53.

45. Harlan JM, Killen PD, Harker LA, Striker GE. of Washington. 1981;68(July):1394– 403.

46. Fujie K, Shinguh Y, Inamura N, Yasumitsu R, Okamoto M, Okuhara M. Release of neutrophil elastase and its role in tissue injury in acute inflammation: Effect of the elastase inhibitor, FR134043. Eur J Pharmacol. 1999;374(1):117–25.

47. Owen CA, Campbell EJ. The cell biology of leukocyte-mediated proteolysis. 1999;65(February):137–50.

48. Thompson RC, Ohlsson K. Isolation, properties, and complete amino acid sequence of human secretory leukocyte protease inhibitor, a potent inhibitor of leukocyte elastase. Proc Natl Acad Sci U S A [Internet]. 1986 [cited 2021 Apr 13];83(18):6692–6. Available from: https://pubmed.ncbi.nlm.nih.gov/3462719/

49. Taggart CC, Cryan SA, Weldon S, Gibbons A, Greene CM, Kelly E, et al. Secretory leucoprotease inhibitor binds to NF-κB binding sites in monocytes and inhibits p65 binding. Journal of Experimental Medicine [Internet]. 2005 Dec 19 [cited 2021 Apr 13];202(12):1659–68. Available from: https://pubmed.ncbi.nlm.nih.gov/16352738/

50. Ward PA, Lentsch AB. Endogenous regulation of the acute inflammatory response. 2002;225–8.

51. Nugteren S, Simons-Oosterhuis Y, Menckeberg CL, Hulleman-van Haaften DH, Lindenbergh-Kortleve DJ, Samsom JN. Endogenous secretory leukocyte protease inhibitor inhibits microbial-induced monocyte activation. Eur J Immunol. 2023;53(2):1–13.

52. Schilling JD, Mulvey MA, Vincent CD, Lorenz RG, Hultgren SJ. Bacterial Invasion Augments Epithelial Cytokine Responses to Escherichia coli Through a Lipopolysaccharide-Dependent Mechanism . The Journal of Immunology [Internet]. 2001 Jan 15 [cited 2021 Apr 13];166(2):1148–55. Available from: https://pubmed.ncbi.nlm.nih.gov/11145696/

53. Wu XR, Sun TT, Medina JJ. In vitro binding of type 1-fimbriated Escherichia coli to uroplakins Ia and Ib: Relation to urinary tract infections. Proc Natl Acad Sci U S A [Internet]. 1996 Sep 3 [cited 2021 Apr 13];93(18):9630–5. Available from: https://pubmed.ncbi.nlm.nih.gov/8790381/

54. Justice SS, Lauer SR, Hultgren SJ, Hunstad DA. Maturation of intracellular Escherichia coli communities requires SurA. Infect Immun. 2006;74(8):4793–800.

55. Wiedow O, Harder J, Bartels J, Streit V, Christophers E. Antileukoprotease in human skin: An antibiotic peptide constitutively produced by keratinocytes. Biochem Biophys Res Commun. 1998;248(3):904–9.

56. Chromek M, Arvidsson I, Karpman D. The Antimicrobial Peptide Cathelicidin Protects Mice from Escherichia coli O157:H7-Mediated Disease. PLoS One. 2012;7(10):1–8.

57. Hopkins WJ, Gendron-Fitzpatrick A, Balish E, Uehling DT. Time course and host responses to Escherichia coli urinary tract infection in genetically distinct mouse strains. Infect Immun [Internet]. 1998 [cited 2021 Apr 13];66(6):2798–802. Available from: https://pubmed.ncbi.nlm.nih.gov/9596750/

58. Reardon C, Lechmann M, Brüstle A, Gareau MG, Shuman N, Philpott D, et al. Thymic Stromal Lymphopoetin-Induced Expression of the Endogenous Inhibitory Enzyme SLPI Mediates Recovery from Colonic Inflammation. Immunity [Internet]. 2011 Aug 26 [cited 2021 Apr 13];35(2):223–35. Available from: https://pubmed.ncbi.nlm.nih.gov/21820333/

59. Marino R, Thuraisingam T, Camateros P, Kanagaratham C, Xu YZ, Henri J, et al. Secretory Leukocyte Protease Inhibitor Plays an Important Role in the Regulation of Allergic Asthma in Mice. The Journal of Immunology [Internet]. 2011 Apr 1 [cited 2021 Apr 13];186(7):4433–42. Available from: https://pubmed.ncbi.nlm.nih.gov/21335488/

60. Jaeger N, McDonough RT, Rosen AL, Hernandez-Leyva A, Wilson NG, Lint MA, et al. Airway Microbiota-Host Interactions Regulate Secretory Leukocyte Protease Inhibitor Levels and Influence Allergic Airway Inflammation. Cell Rep [Internet]. 2020 Nov 3 [cited 2021 Apr 13];33(5). Available from: https://pubmed.ncbi.nlm.nih.gov/33147448/

61. Conover MS, Flores-Mireles AL, Hibbing ME, Dodson K, Hultgren SJ. Establishment and characterization of UTI and CAUTI in a mouse model. Journal of Visualized Experiments. 2015;2015(100):1–12.

62. Lentsch AB, Jordan JA, Czermak BJ, Diehl KM, Younkin EM, Sarma V, et al. Inhibition of NF-κB activation and augmentation of IκBβ by secretory leukocyte protease inhibitor during lung inflammation. American Journal of Pathology. 1999;154(1):239–47.

63. Skrzeczynska-Moncznik J, Zabieglo K, Osiecka O, Morytko A, Brzoza P, Drozdz L, et al. Differences in Staining for Neutrophil Elastase and its Controlling Inhibitor SLPI Reveal Heterogeneity among Neutrophils in Psoriasis. Journal of Investigative Dermatology. 2020;140(7):1371–1378.e3.

64. Ozaka S, Sonoda A, Ariki S, Kamiyama N, Hidano S, Sachi N, et al. Protease inhibitory activity of secretory leukocyte protease inhibitor ameliorates murine experimental colitis by protecting the intestinal epithelial barrier. Genes to Cells. 2021;26(10):807–22.

65. Hagio T, Nakao S, Matsuoka H, Matsumoto S, Kawabata K, Ohno H. Inhibition of neutrophil elastase activity attenuates complement-mediated lung injury in the hamster. Eur J Pharmacol. 2001;426(1–2):131–8.

66. Chul Hee Lee, Igarashi Y, Hohman RJ, Kaulbach H, White M V, Kaliner MA. Distribution of secretory leukoprotease inhibitor in the human nasal airway. American Review of Respiratory Disease. 1993;147(3):710–6.

67. Abe T, Kobayashi N, Yoshimura K, Trapnell BC, Kim H, Hubbard RC, et al. Expression of the secretory leukoprotease inhibitor gene in epithelial cells. Journal of Clinical Investigation. 1991;87(6):2207–15.

68. Jin FY, Nathan C, Radzioch D, Ding A. Secretory leukocyte protease inhibitor: A macrophage product induced by and antagonistic to bacterial lipopolysaccharide. Cell. 1997;88(3):417–26.

69. Zabieglo K, Majewski P, Majchrzak-Gorecka M, Wlodarczyk A, Grygier B, Zegar A, et al. The inhibitory effect of secretory leukocyte protease inhibitor (SLPI) on formation of neutrophil extracellular traps. J Leukoc Biol [Internet]. 2015 Jul [cited 2021 Apr 13];98(1):99–106. Available from: https://pubmed.ncbi.nlm.nih.gov/25917460/

70. Osiecka O, Skrzeczynska-Moncznik J, Morytko A, Mazur A, Majewski P, Bilska B, et al. Secretory Leukocyte Protease Inhibitor Is Present in Circulating and Tissue-Recruited Human Eosinophils and Regulates Their Migratory Function. Front Immunol. 2022;12(January):1–11.

71. Layland LE, Mages J, Loddenkemper C, Hoerauf A, Wagner H, Lang R, et al. Pronounced Phenotype in Activated Regulatory T Cells during a Chronic Helminth Infection. The Journal of Immunology [Internet]. 2010 Jan 15 [cited 2023 Jan 12];184(2):713–24. Available from: https://journals.aai.org/jimmunol/article/184/2/713/111881/Pronounced-Phenotype-in-Activated-Regulatory-T

72. Li J, Peet GW, Balzarano D, Li X, Massa P, Barton RW, et al. Novel NEMO/IκB Kinase and NF-κB Target Genes at the Pre-B to Immature B Cell Transition. Journal of Biological Chemistry [Internet]. 2001;276(21):18579–90. Available from: 10.1074/jbc.M100846200

73. Birrer P, McElvaney NG, Gillissen A, Hoyt RF, Bloedow DC, Hubbard RC, et al. Intravenous recombinant secretory leukoprotease inhibitor augments antineutrophil elastase defense. J Appl Physiol [Internet]. 1992 [cited 2021 Apr 13];73(1):317–23. Available from: https://pubmed.ncbi.nlm.nih.gov/1354669/

74. Bergenfeldt M, Björk P, Ohlsson K. The elimination of secretory leukocyte protease inhibitor (SLPI) after intravenous injection in dog and man. Scand J Clin Lab Invest [Internet]. 1990 [cited 2021 Apr 13];50(7):729–37. Available from: https://pubmed.ncbi.nlm.nih.gov/2293334/

75. Mowbray C, Tan A, Vallée M, Fisher H, Chadwick T, Brennand C, et al. Multidrug-resistant Uro-associated Escherichia coli Populations and Recurrent Urinary Tract Infections in Patients Performing Clean Intermittent Self-catheterisation. Eur Urol Open Sci. 2022 Mar 1;37:90–8.

76. Schiwon M, Weisheit C, Franken L, Gutweiler S, Dixit A, Meyer-Schwesinger C, et al. Crosstalk between sentinel and helper macrophages permits neutrophil migration into infected uroepithelium. Cell [Internet]. 2014 Jan 30 [cited 2021 Apr 13];156(3):456–68. Available from: https://pubmed.ncbi.nlm.nih.gov/24485454/

77. Fernie-King BA, Seilly DJ, Davies A, Lachmann PJ. Streptococcal inhibitor of complement inhibits two additional components of the mucosal innate immune system: Secretory leukocyte proteinase inhibitor and lysozyme. Infect Immun. 2002;70(9):4908–16.

78. Osbourn M, Rodgers AM, Dubois A V., Small DM, Humphries F, Delagic N, et al. Secretory Leucoprotease Inhibitor (SLPI) Promotes Survival during Acute Pseudomonas aeruginosa Infection by Suppression of Inflammation Rather Than Microbial Killing. Biomolecules. 2022;12(12).

79. Parameswaran GI, Wrona CT, Murphy TF, Sethi S. Moraxella catarrhalis acquisition, airway inflammation and protease-antiprotease balance in chronic obstructive pulmonary disease. BMC Infect Dis. 2009;9:1–10.

80. Parameswaran GI, Sethi S, Murphy TF. Effects of bacterial infection on airway antimicrobial peptides and proteins in COPD. Chest. 2011;140(3):611–7.

81. Vanden Berghe T, Hulpiau P, Martens L, Vandenbroucke RE, Van Wonterghem E, Perry SW, et al. PasSenger Mutations Confound Interpretation Of All Genetically Modified Congenic Mice. Immunity. 2015;43(1):200–9.

82. Morimoto A, Kikuta J, Nishikawa K, Sudo T, Uenaka M, Furuya M, et al. SLPI is a critical mediator that controls PTH-induced bone formation. Nat Commun. 2021;12(1):1–14.

83. Datsenko KA, Wanner BL. One-step inactivation of chromosomal genes in Escherichia coli K-12 using PCR products. Proc Natl Acad Sci U S A. 2000;97(12):6640–5.

84. Murphy KC, Campellone KG. Lambda Red-mediated recombinogenic engineering of enterohemorrhagic and enteropathogenic E. coli. BMC Mol Biol. 2003;4:1–12.

85. Livak KJ, Schmittgen TD. Analysis of relative gene expression data using real-time quantitative PCR and the 2-ΔΔCT method. Methods. 2001;25(4):402–8.

86. Patro R, Duggal G, Love MI, Irizarry RA, Kingsford C. Salmon provides fast and bias-aware quantification of transcript expression. Nat Methods. 2017;14(4):417– 9.

87. Love MI, Huber W, Anders S. Moderated estimation of fold change and dispersion for RNA-seq data with DESeq2. Genome Biol. 2014;15(12):1–21.

88. Durinck S, Spellman PT, Birney E, Huber W. Mapping identifiers for the integration of genomic datasets with the R/ Bioconductor package biomaRt. Nat Protoc. 2009;4(8):1184–91.

89. Korotkevich G, Sukhov V. Fast gene set enrichment analysis. 2016;1–29.

